## Supplemental for "Structural basis of Spliced Leader RNA recognition by the *Trypanosoma brucei* cap-binding complex"

**Figure S1: (A) Purity of *Tb*CBC complex samples.** A Coomassie-stained 12% SDS-PAGE of the purified protein and protein complexes used in the study, representative gel of more than three replicates. **(B)** Mass photometry histograms of the samples.

**Figure S2: EM processing scheme of *Tb*CBC-tetramer. (A)** Representative micrograph and schematic of the processing steps used. Particles from classes in blue were pooled for subsequent steps, and particles from classes in grey were discarded. **(B)** Final map from RELION 3D refinement post-processed by DeepEMhancer. **(C)** Local resolution. **(D)** Angular distribution map of particle orientation of the final refinement step in RELION. **(E)** Final resolution of the reconstruction, as calculated by RELION Gold Standard Fourier Shell Correlation, with a resolution of 2.4 Å when adopting a cutoff of 0.143.

**Figure S3: Architecture of *Tb*CBC. (A) Small-angle** X-ray scattering (SAXS) of *Tb*CBC dimer and tetramer. **(B)** The CBP20-CBP110 interface. Surface representation of *Tb*CBC-tetramer colored as in main Figure 1 (TbCBP110 green, *Tb*CBP20 yellow, *Tb*CBP30 pink) and surface representation of individual *Tb*CBP110 and *Tb*CBP20, colored by surface electrostatic charge as determined by ChimeraX's coulombic tool.

**Figure S4: Synthesis of *T. brucei* cap4 ligand. (A, B, C)** Schematics of the individual steps involved in the preparation of cap4. **(D)** Example HPLC profile of $m^7Gppp^{m6,6}A_mpA_mpCp^{m3}U_mpA$.

**Figure S5: EM processing scheme of *Tb*CBC-tetramer. (A)** Representative micrograph and schematic of the processing steps used. Particles from classes in blue were pooled for subsequent steps, and particles from classes in grey were discarded. **(B)** Final map from cryosparc homogenous refinement and sharpened by DeepEMhancer. **(C)** Local resolution. **(D)** Angular distribution map of particle orientation of the final refinement step in CryoSPARC. **(E)** The final resolution of the reconstruction, as calculated by CryoSPARC Gold Standard Fourier Shell Correlation, with a resolution of 2.8 Å when adopting a cutoff of 0.143.

**Figure S6: Comparison between *Tb*CBC-tetramer and *Tb*CBC-trimer models, and *T. brucei* and human CBP20. (A)** Superimposition of *Tb*CBC-tetramer and *Tb*CBC-trimer in two orientations in cartoon representation. **(B)** Zoom into the differences between *Tb*CBC-tetramer and *Tb*CBC-trimer models comprising the C-terminal domain of *Tb*CBP20 and regions of *Tb*CBP30: *Tb*CBC-tetramer in **(B)** and *Tb*CBC-trimer in **(C). (D)** Side-by-side comparison of *Tb*CBP20 and *Hs*CBP20 (PDB:1H2T). Colors as in main Figure 1 and 3 (for *Tb*CBP20 domains). **(E)** CryoSPARC final homogeneous refinement trimer cryo-EM map transparent coloured as Figura 3 with cartoon model and m7GTP in stick representation and at level 0.44 (left) and at a very reduced level of 0.22 (right). Unassigned density at the reduced level map in light gray.

**Figure S7: Sequence alignment of *Tb*CBC subunits. (A)** Alignment of *Tb*CBP20 with the CBP20 of other trypanosomatids and to CBP20 of selected Opisthokonts. Black triangles highlight residues involved in cap binding. Black circles highlight kinetoplast-conserved residues involved in the interaction with cap4. Red circles highlight kinetoplast-conserved residues involved in the interaction with *Tb*CBP30 **(B)** Alignment of *Tb*CBP30 with the CBP30 of other trypanosomatids. Regions involved with interaction with *Tb*CBP20 and *Tb*CBP110 are highlighted in yellow and green, respectively, with residues mutated in this study highlighted with circles. The region used for the *Tb*CBP66* fusion construct is highlighted in purple. **(C)** Alignment of *Tb*CBP66 with the CBP66 of other trypanosomatids. Residues mutated in this study are highlighted with a red star. **(D)** Alignment of *Tb*CBP110 with the CBP110 of other trypanosomatids. Secondary structure annotations based on our structure for CBP20 and CBP110 and on AlphaFold DB prediction for CBP30 and CBP66

**Figure S8: Synthesis of *T.brucei* cap0 SL RNA (40 nt) and cap4 SL RNA (40 nt). (A)** Schematics of the individual steps involved in the preparation of cap4 RNA. The cap0 analog was prepared following the same procedure. **(B)** Left traces: Solid phase synthesis of short Gppp-RNA (steps 1-5). Reaction control of the individual steps was based on small portions of support-bound RNAs (I to III) that were individually deprotected and analyzed by AE HPLC. Right traces: Guanine N7 methylation of Gppp-RNA (step 6). HPLC profiles show the methylation reaction at the start (HPLC-purified oligomer III) and after 1 h; inset shows purified product of methylation (IV). **(C)** Example of enzymatic ligation of cap4 RNA (18 nt) and chemically synthesized 5'-p RNA (22 nt) using T4 DNA ligase and DNA splint (23 nt). The left HPLC profiles show sample composition at the start and after completion of the reaction (3 h). The middle traces show the HPLC-purified ligation products (V) that were further characterized by mass spectrometry (right traces).

**Figure S9: *Tb*CBC30 interactions. (A)** AlphaFold2 model of *Tb*CBP30 and corresponding predicted alignment error (PAE) plot. The *Tb*CBP20 interaction region is colored in pink, and the helical region used to create the *Tb*CBP66* fusion construct is colored in blue. **(B)** Co-immunoprecipitation assay of co-expressed *Tb*CBP20, *Tb*CBP110, *Tb*CBP66 with mCherry-*Tb*CBP30 mutants and Western blot detection (compare to main Figure 4C). **(C)** Comparison of the NELF-E binding on *human* CBP20-CBP80 dimer (left) with the *Tb*CBP30 peptide binding on *Tb*CBP20-*Tb*CBP110 dimer (middle). Right panel: only the binding peptides are shown. **(D)** Comparison of binding of arginine R35 (*Tb*CBP30) with the binding or arginines in the same position of the human co-factors ARS2 and NELF-E on the human CBC (PDB: 5OO6 and 5OOB).

**Figure S10:** Microscale thermophoresis SDS-denaturation (SD) test, used to validate the approach of initial fluorescence variation for the dissociation constant estimation. The top panel represents the capillary scan, and the bottom represents the plotted data, with an experimental run in green and the SD test in red.

#### Figure S1

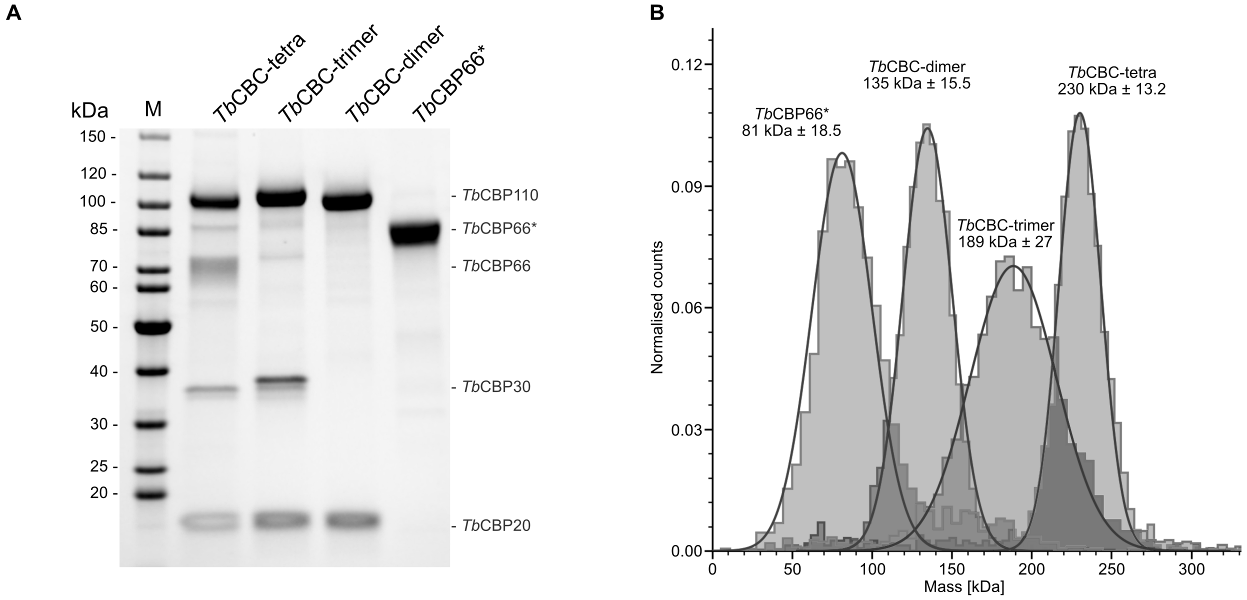
A
kDa
M
TbCBC-tetra
TbCBC-trimer
TbCBC-dimer
TbCBP66*
150 -
120 -
100 -
85 -
70 -
60 -
50 -
40 -
30 -
25 -
20 -
- TbCBP110
- TbCBP66*
- TbCBP66
- TbCBP30
- TbCBP20
B
TbCBP66*
81 kDa ± 18.5
TbCBC-dimer
135 kDa ± 15.5
TbCBC-trimer
189 kDa ± 27
TbCBC-tetra
230 kDa ± 13.2
Normalised counts
0.12
0.09
0.06
0.03
0.00
0
50
100
150
200
250
300
Mass [kDa]

**Figure S2**

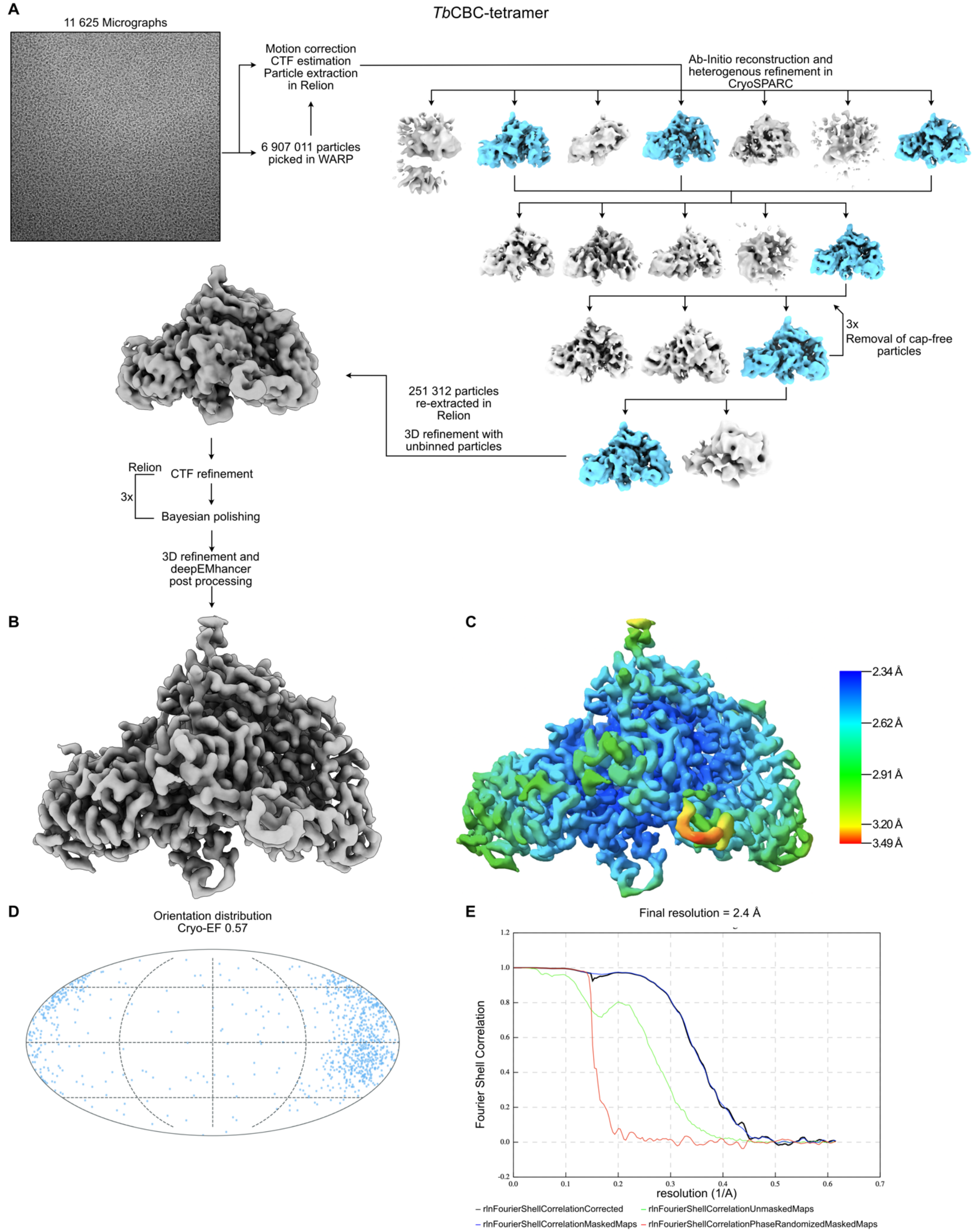
A
*Tb*CBC-tetramer
11 625 Micrographs
Motion correction
CTF estimation
Particle extraction
in Relion
6 907 011 particles
picked in WARP
Ab-Initio reconstruction and
heterogenous refinement in
CryoSPARC
3x
Removal of cap-free
particles
251 312 particles
re-extracted in
Relion
3D refinement with
unbinned particles
Relion
CTF refinement
3x
Bayesian polishing
3D refinement and
deepEMhancer
post processing
B
C
2.34 Å
2.62 Å
2.91 Å
3.20 Å
3.49 Å
D
Orientation distribution
Cryo-EF 0.57
E
Final resolution = 2.4 Å
Fourier Shell Correlation
resolution (1/A)
rlnFourierShellCorrelationCorrected
rlnFourierShellCorrelationUnmaskedMaps
rlnFourierShellCorrelationMaskedMaps
rlnFourierShellCorrelationPhaseRandomizedMaskedMaps

#### Figure S3

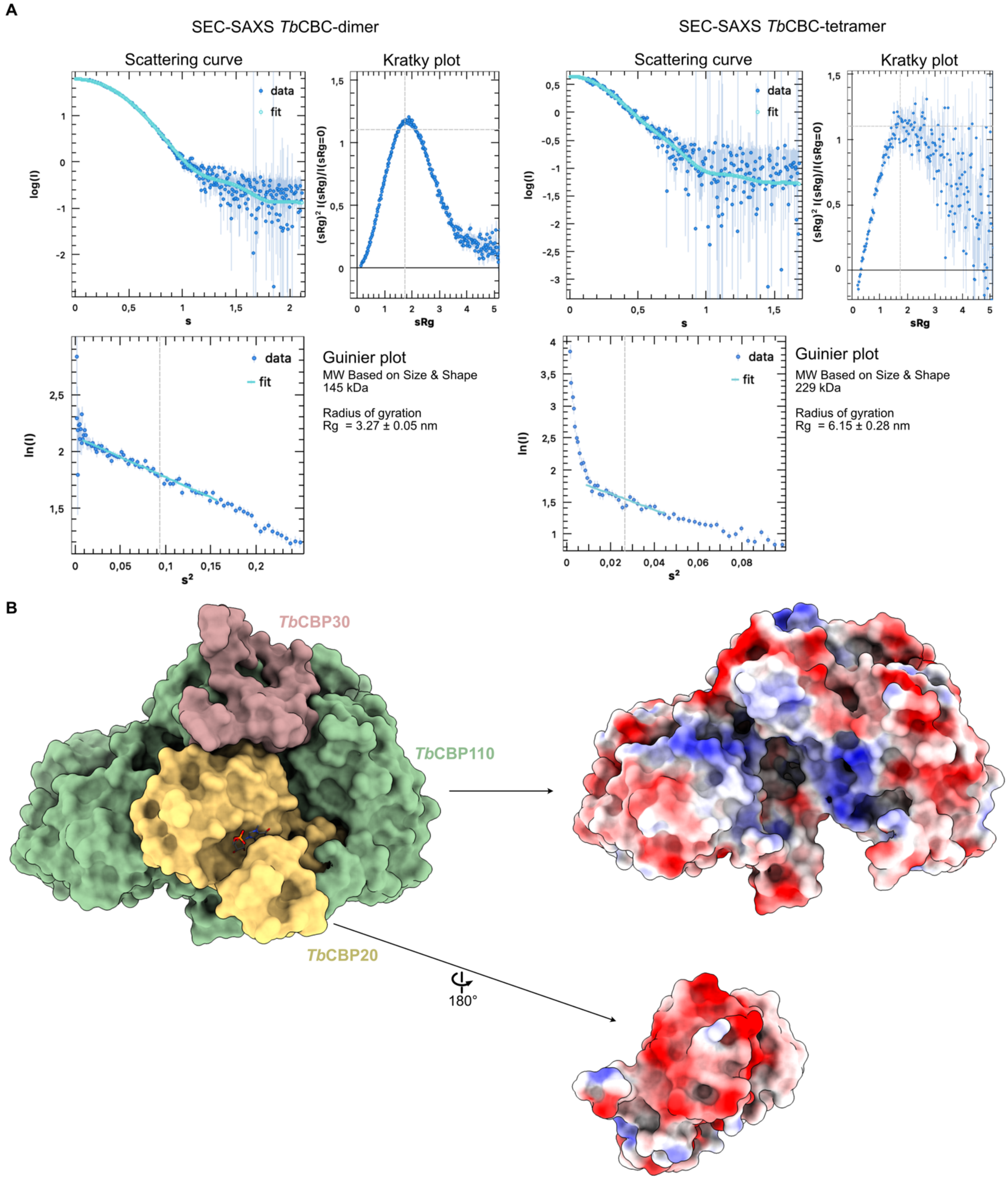
A
SEC-SAXS *Tb*CBC-dimer
Scattering curve
Kratky plot
data
fit
log(I)
s
sRg
(sRg)² I(sRg)/I(sRg=0)
ln(I)
s²
Guinier plot
MW Based on Size & Shape
145 kDa
Radius of gyration
Rg = 3.27 ± 0.05 nm
SEC-SAXS *Tb*CBC-tetramer
Scattering curve
Kratky plot
Guinier plot
MW Based on Size & Shape
229 kDa
Radius of gyration
Rg = 6.15 ± 0.28 nm
B
*Tb*CBP30
*Tb*CBP110
*Tb*CBP20
180°

**Figure S4**

**A**

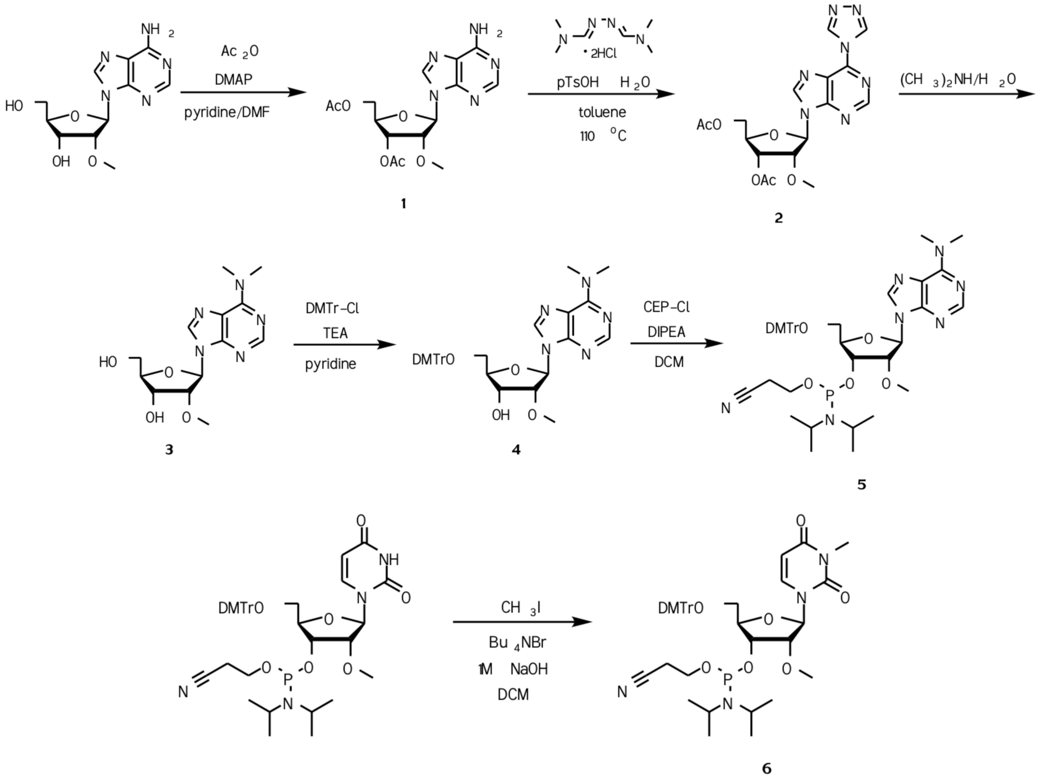


**B**

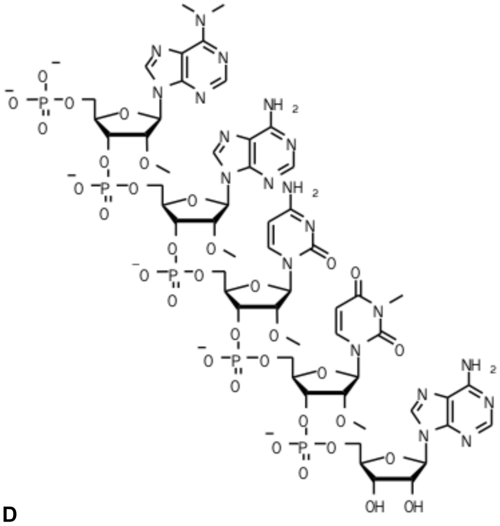


**C**

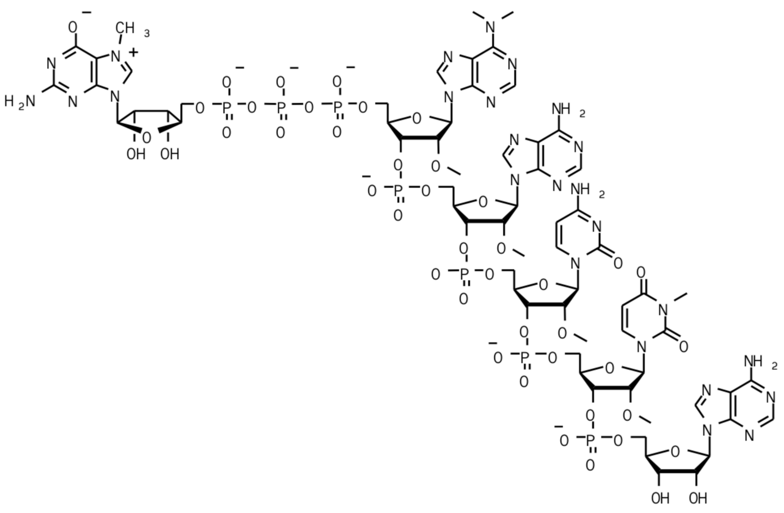


**D**

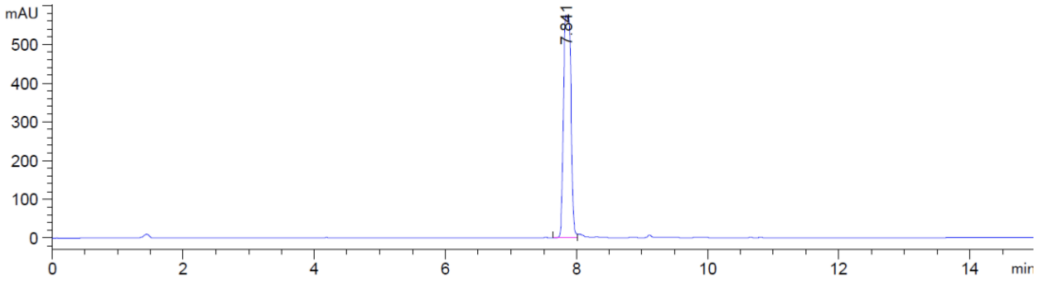

#### Figure S5

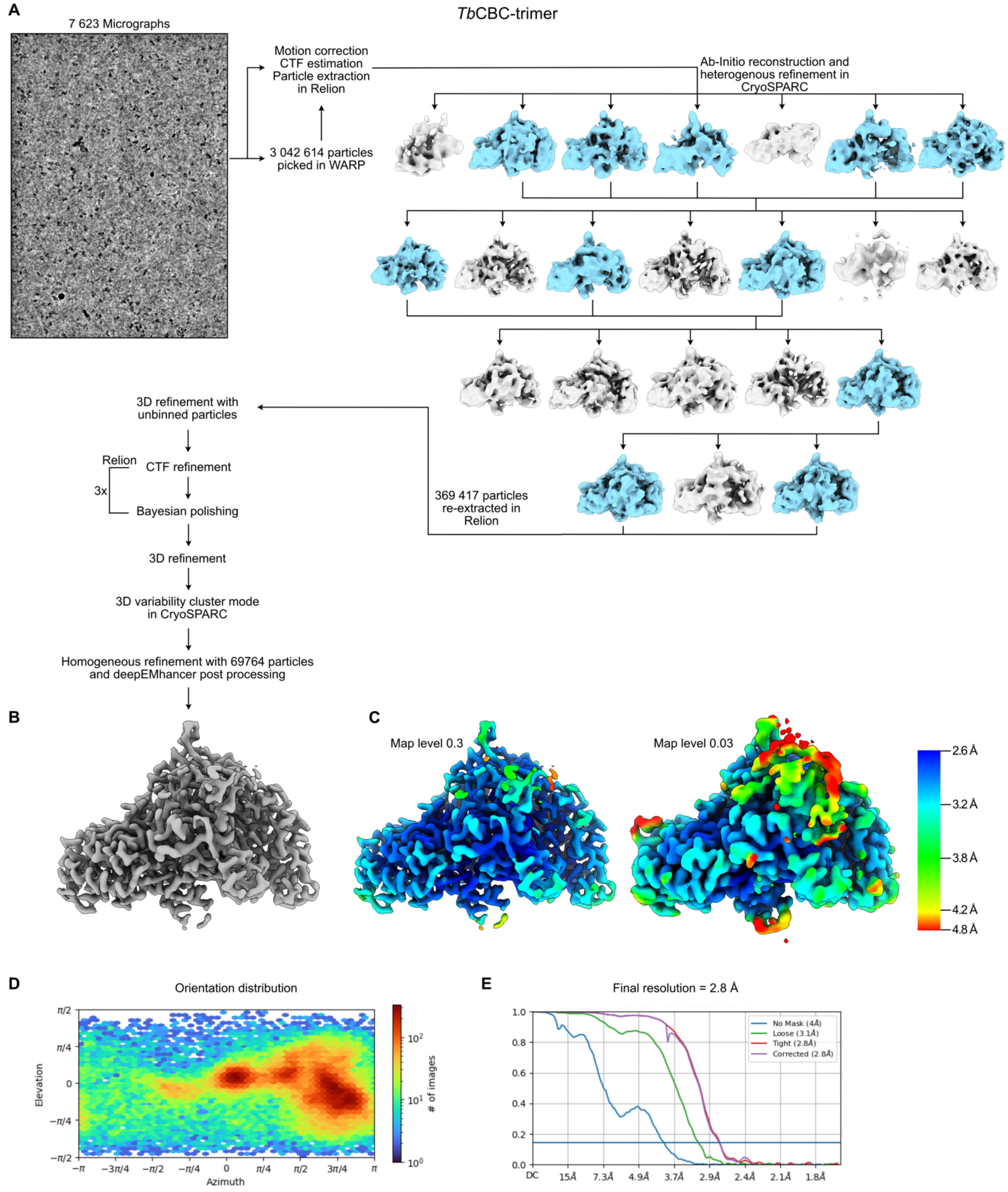

A
7 623 Micrographs
*Tb*CBC-trimer
Motion correction
CTF estimation
Particle extraction
in Relion
3 042 614 particles
picked in WARP
Ab-Initio reconstruction and
heterogenous refinement in
CryoSPARC
3D refinement with
unbinned particles
Relion
CTF refinement
3x
Bayesian polishing
3D refinement
3D variability cluster mode
in CryoSPARC
Homogeneous refinement with 69764 particles
and deepEMhancer post processing
369 417 particles
re-extracted in
Relion
B
C
Map level 0.3
Map level 0.03
2.6 Å
3.2 Å
3.8 Å
4.2 Å
4.8 Å
D
Orientation distribution
Elevation
Azimuth
### of images
E
Final resolution = 2.8 Å
No Mask (4Å)
Loose (3.1Å)
Tight (2.8Å)
Corrected (2.8Å)
DC
15Å
7.3Å
4.9Å
3.7Å
2.9Å
2.4Å
2.1Å
1.8Å

#### Figure S6

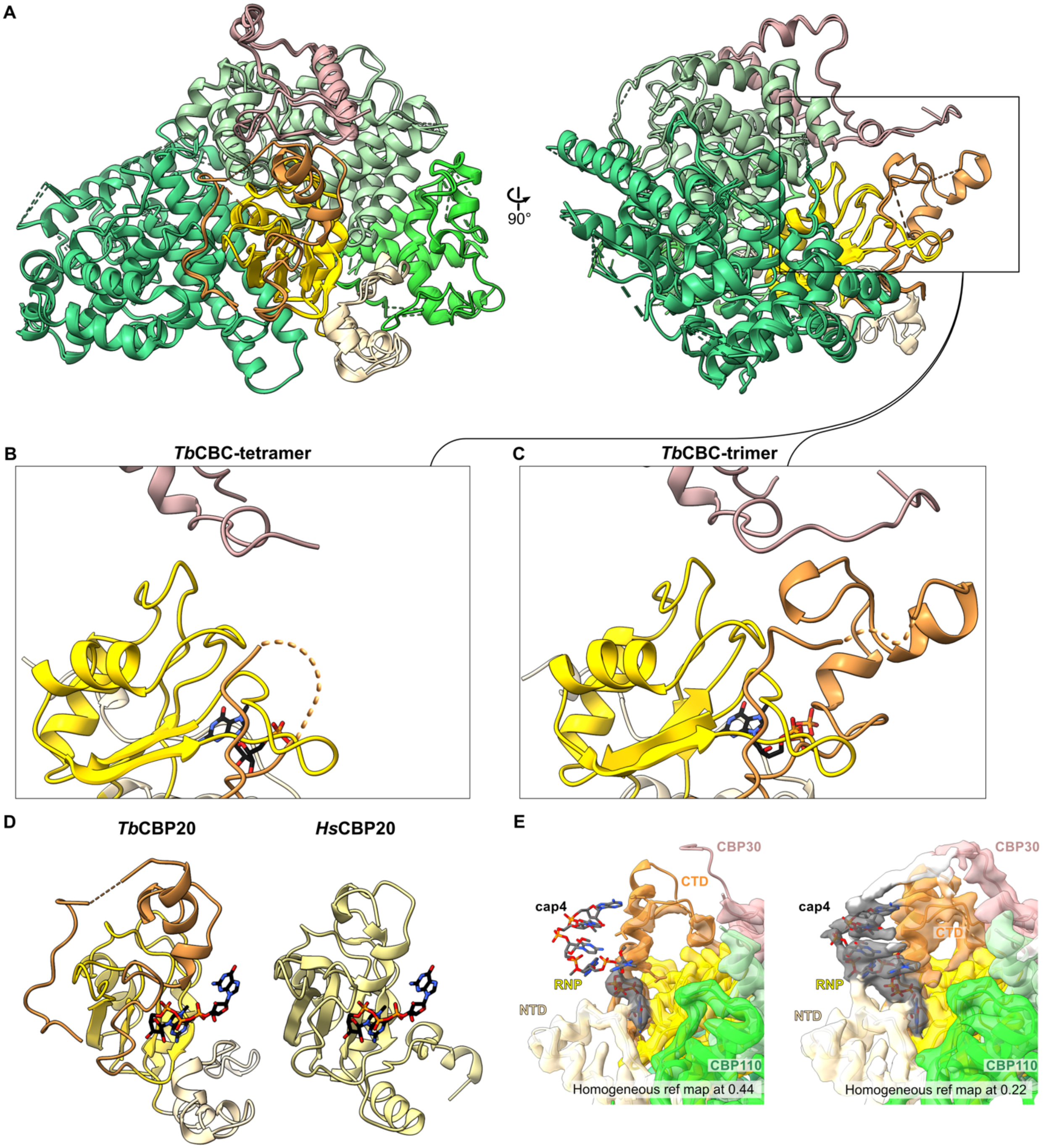

A
90°
B
TbCBC-tetramer
C
TbCBC-trimer
D
TbCBP20
HsCBP20
E
CBP30
CTD
cap4
RNP
NTD
CBP110
Homogeneous ref map at 0.44
CBP30
cap4
CTD
RNP
NTD
CBP110
Homogeneous ref map at 0.22

#### Figure S7

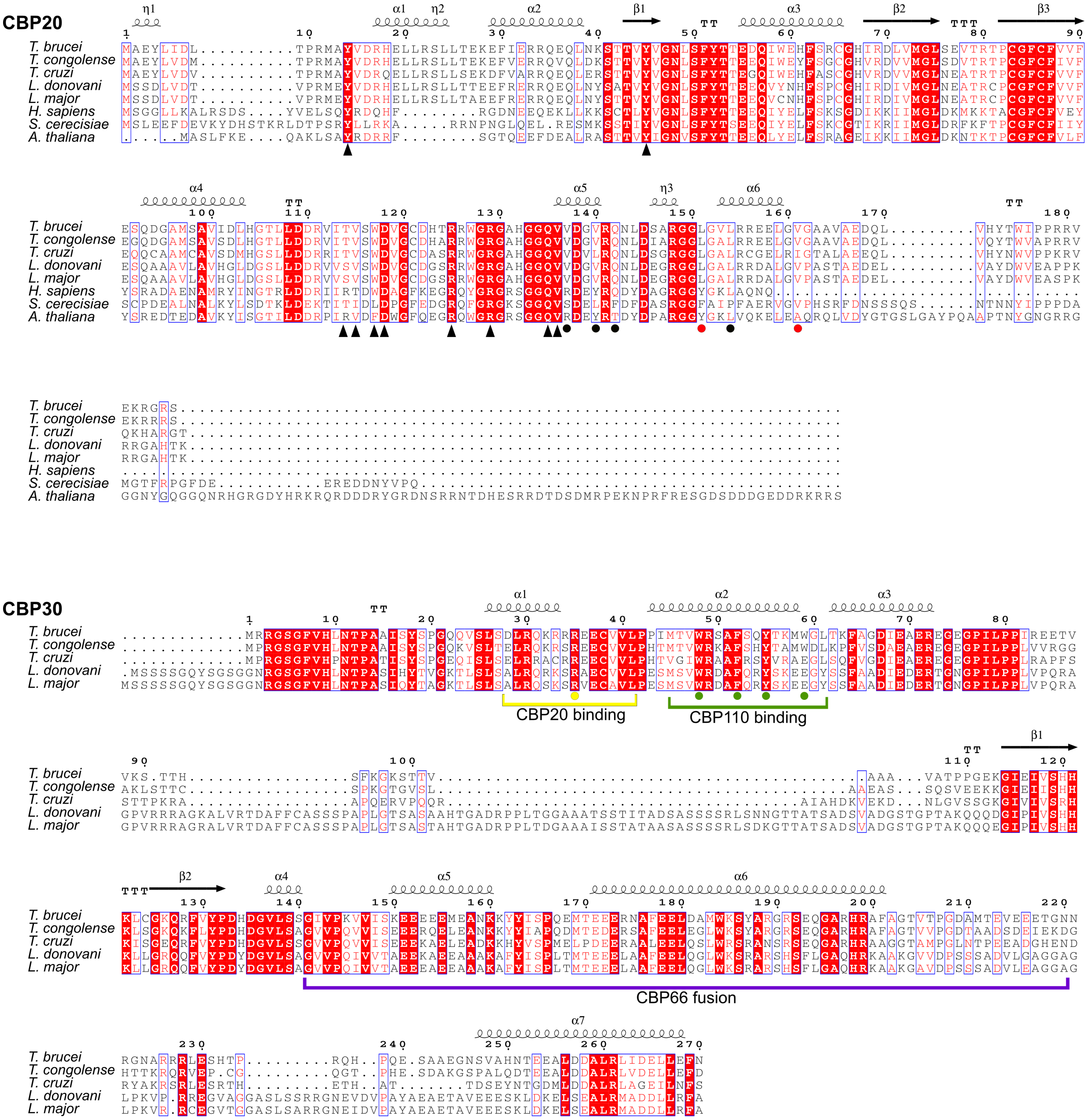

CBP20
T. brucei
T. congolense
T. cruzi
L. donovani
L. major
H. sapiens
S. cerecisiae
A. thaliana
CBP30
CBP20 binding
CBP110 binding
CBP66 fusion

#### Figure S7 (cont.)

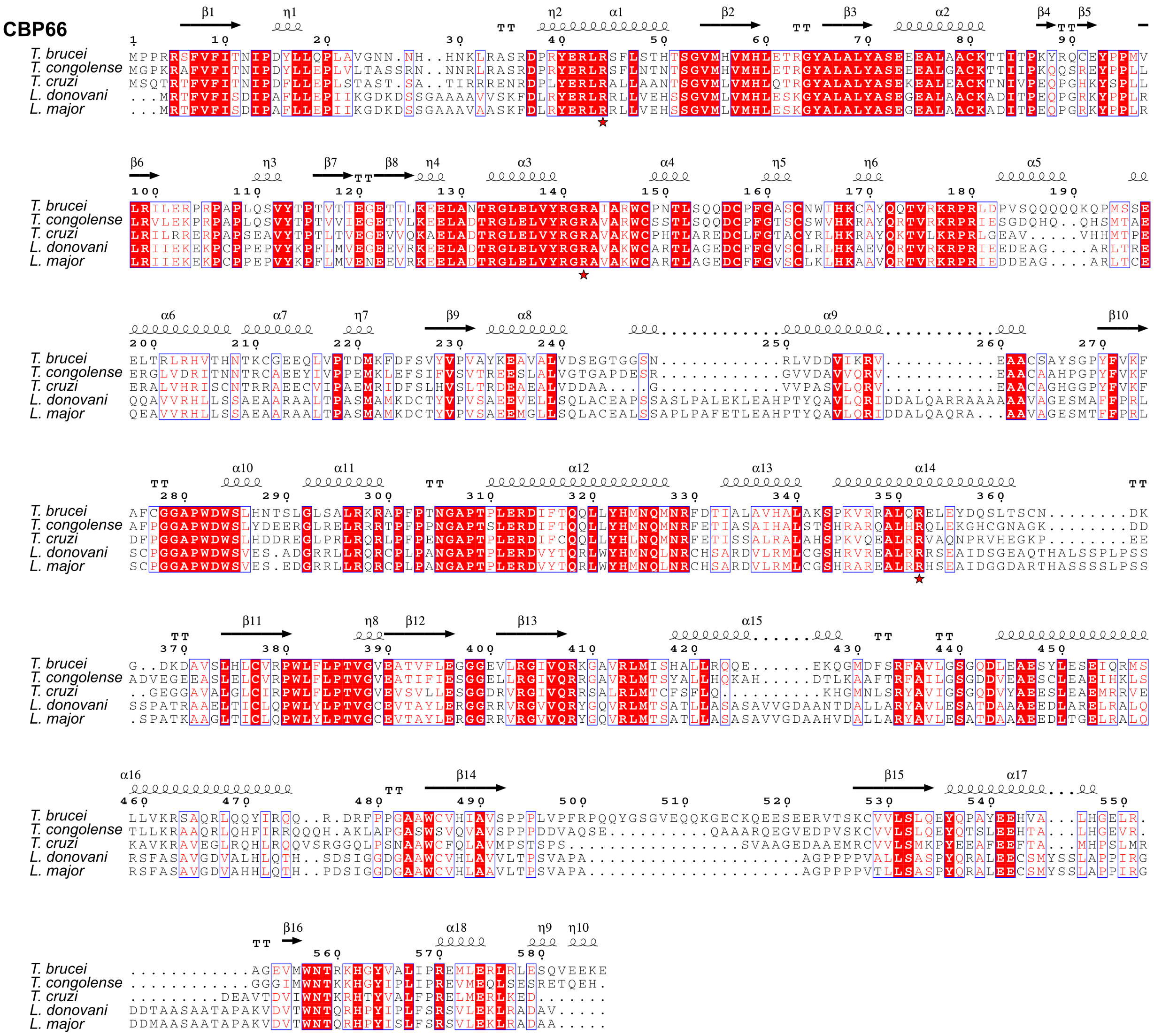

#### Figure S7 (cont.)

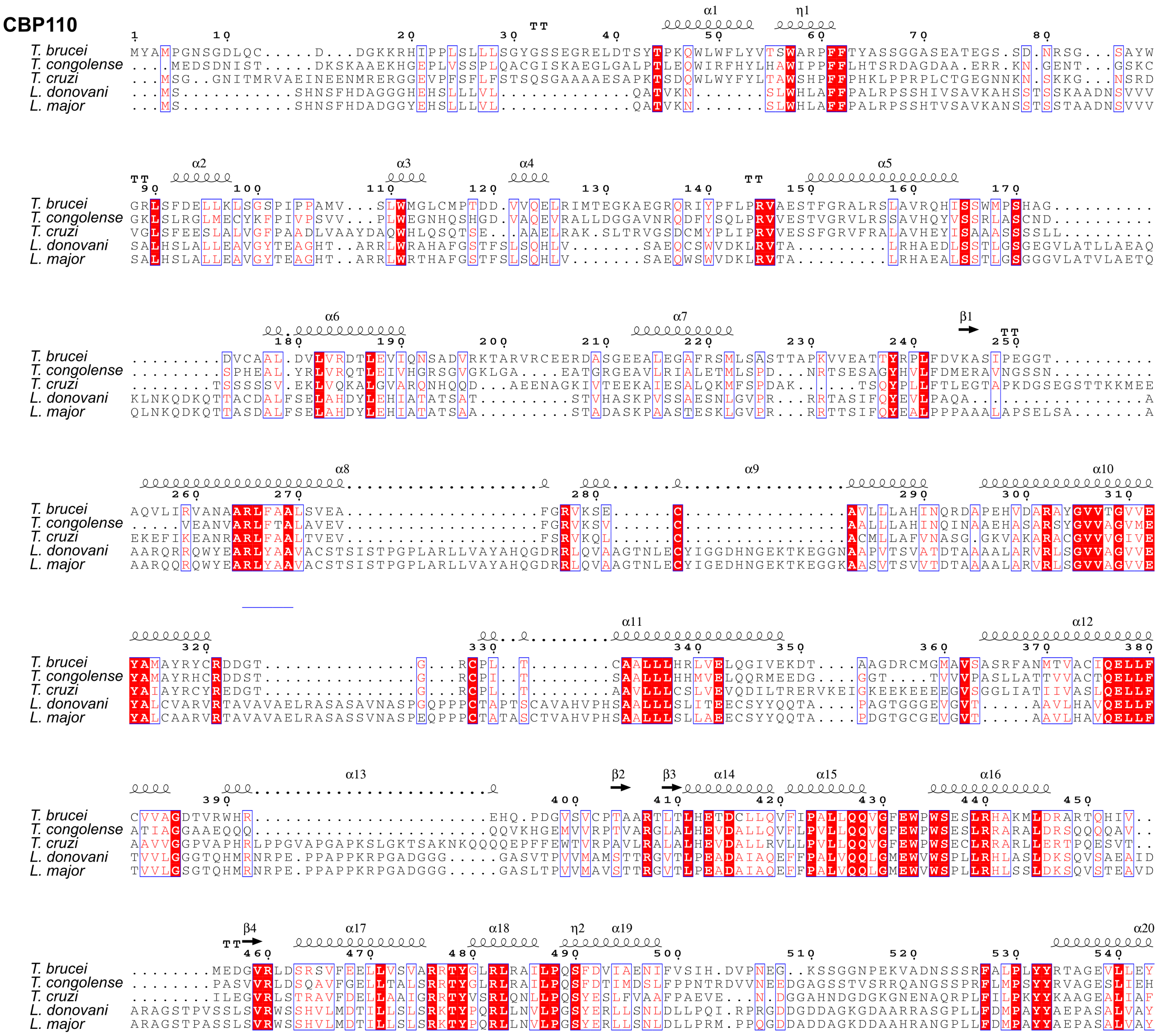

#### Figure S7 (cont.)

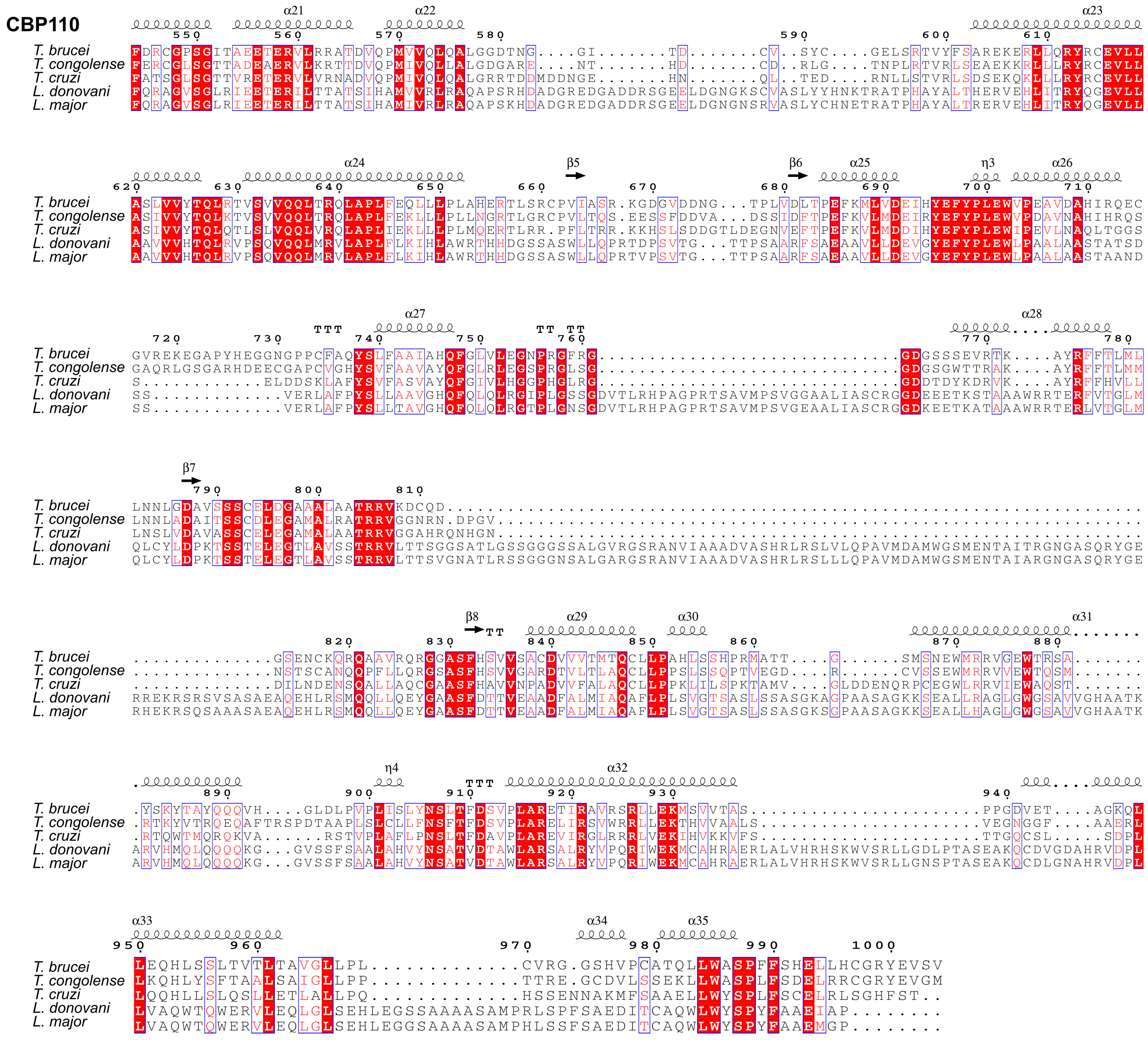

### Figure S8

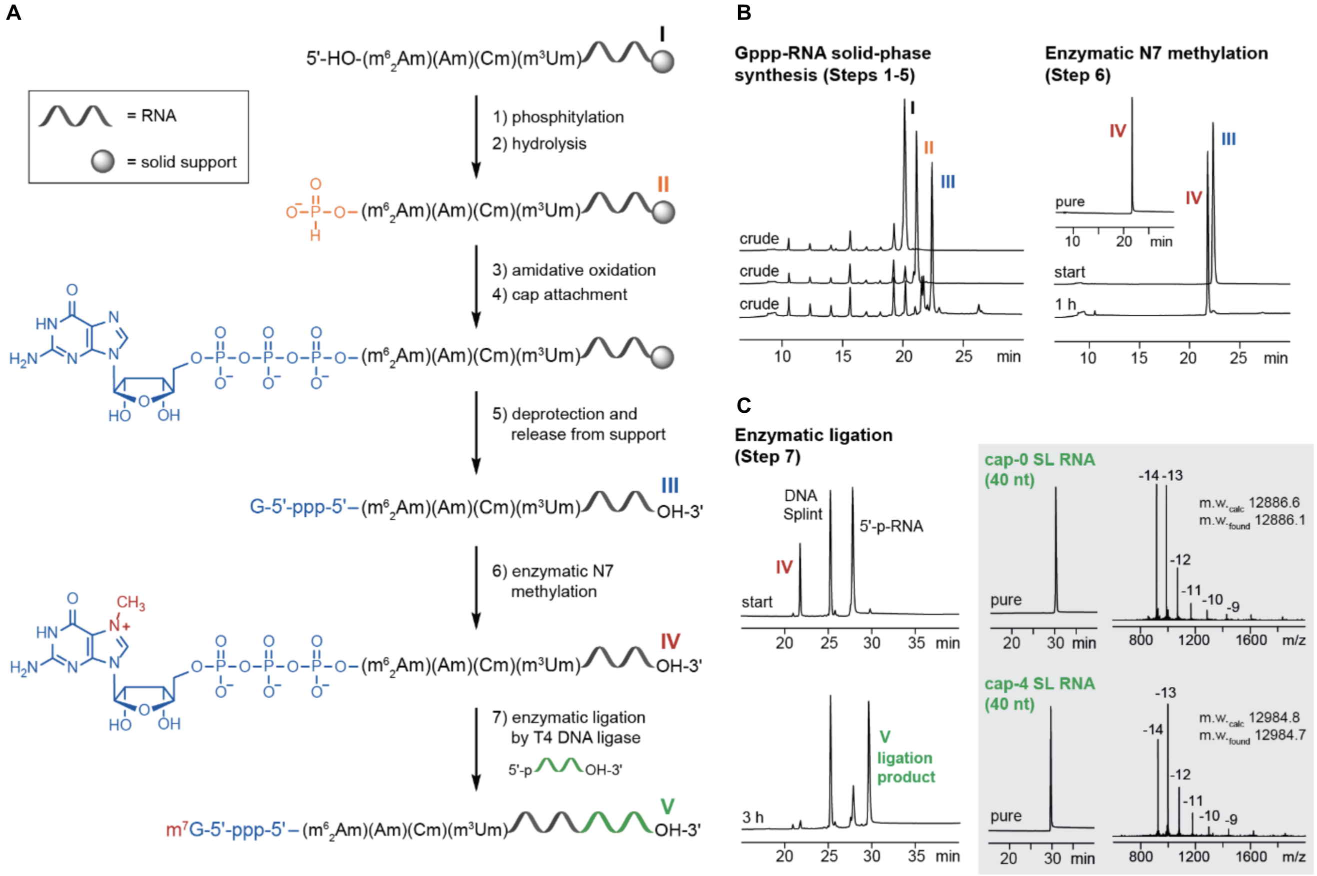
A
= RNA
= solid support
5'-HO-(m62Am)(Am)(Cm)(m3Um)
I
1) phosphitylation
2) hydrolysis
II
3) amidative oxidation
4) cap attachment
5) deprotection and release from support
G-5'-ppp-5'–(m62Am)(Am)(Cm)(m3Um)
OH-3'
III
6) enzymatic N7 methylation
IV
7) enzymatic ligation by T4 DNA ligase
5'-p OH-3'
m7G-5'-ppp-5'–(m62Am)(Am)(Cm)(m3Um)
V
B
Gppp-RNA solid-phase synthesis (Steps 1-5)
crude
crude
crude
Enzymatic N7 methylation (Step 6)
pure
start
1 h
C
Enzymatic ligation (Step 7)
DNA Splint
5'-p-RNA
start
3 h
V ligation product
cap-0 SL RNA (40 nt)
m.w.calc 12886.6
m.w.found 12886.1
cap-4 SL RNA (40 nt)
m.w.calc 12984.8
m.w.found 12984.7
pure
min
m/z

#### Figure S9

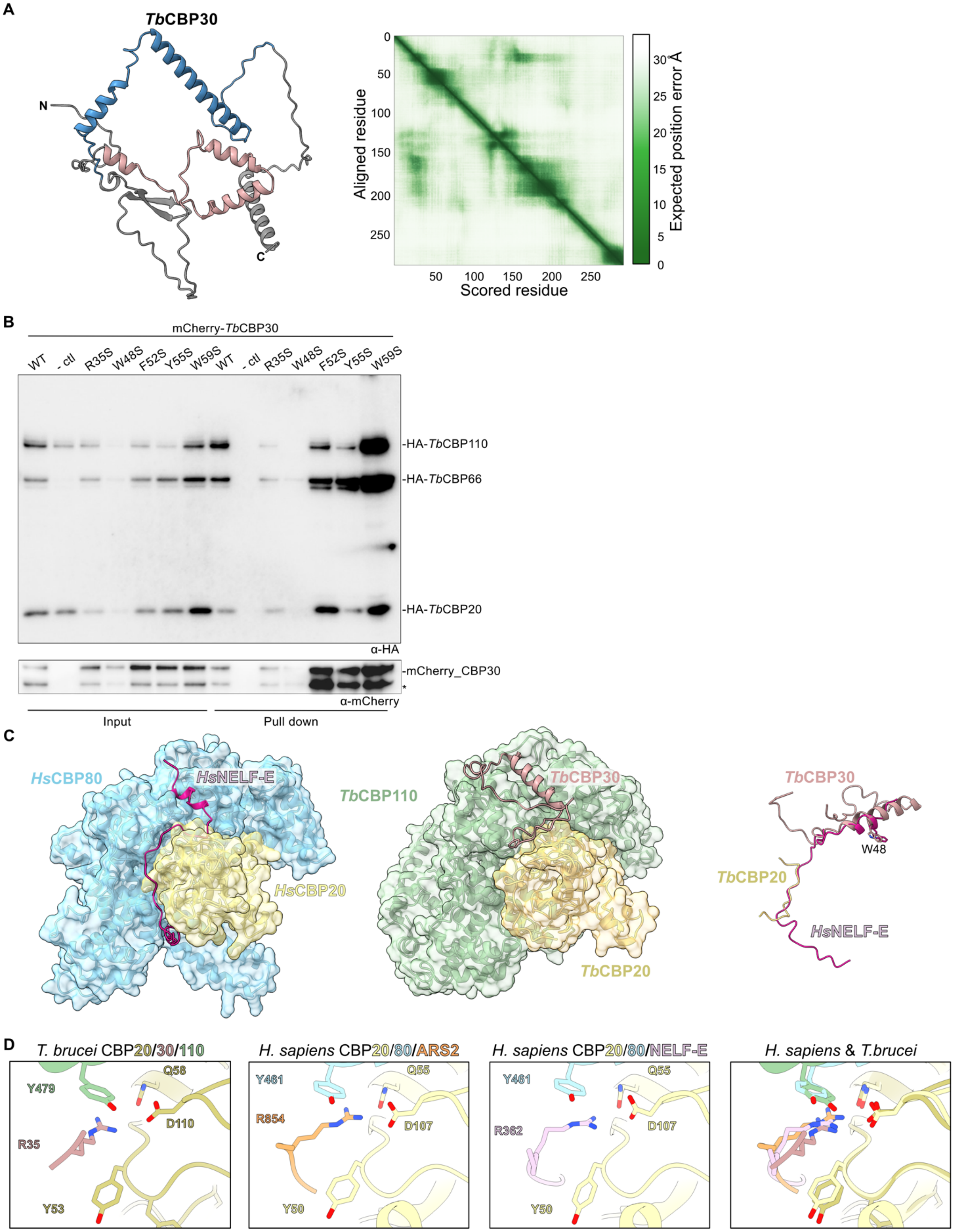

A
TbCBP30
N
C
Aligned residue
Scored residue
Expected position error Å
B
mCherry-TbCBP30
WT
- ctl
R35S
W48S
F52S
Y55S
W59S
-HA-TbCBP110
-HA-TbCBP66
-HA-TbCBP20
α-HA
-mCherry_CBP30
*
α-mCherry
Input
Pull down
C
HsCBP80
HsNELF-E
HsCBP20
TbCBP110
TbCBP30
TbCBP20
W48
D
T. brucei CBP20/30/110
H. sapiens CBP20/80/ARS2
H. sapiens CBP20/80/NELF-E
H. sapiens & T.brucei
Y479
Q58
D110
R35
Y53
Y461
Q55
R854
D107
Y50
R362

**Figure S10**

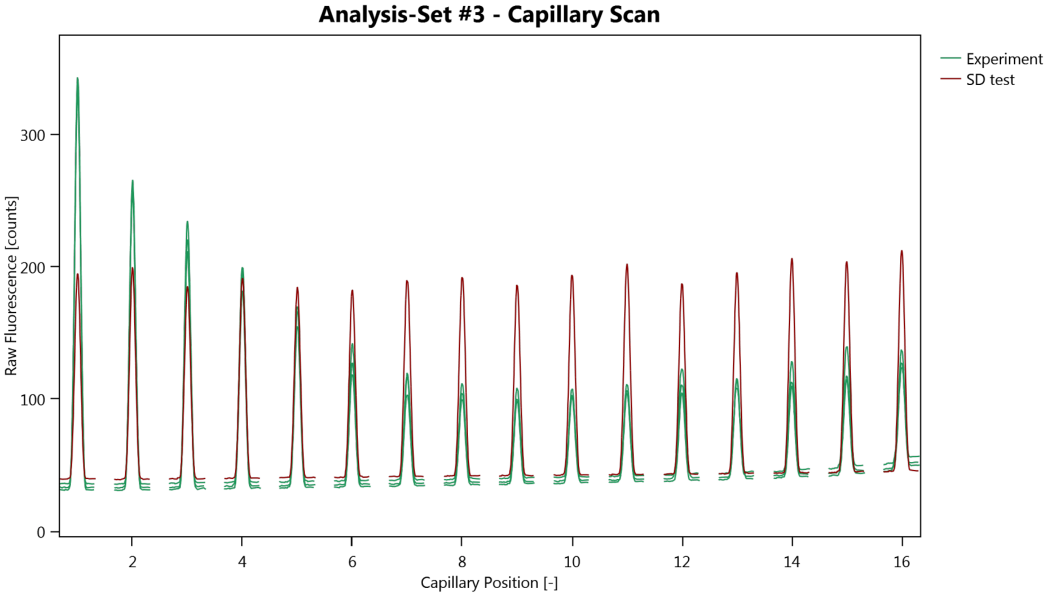

Analysis-Set #3 - Capillary Scan
Experiment
SD test
Raw Fluorescence [counts]
0
100
200
300
2
4
6
8
10
12
14
16
Capillary Position [-]


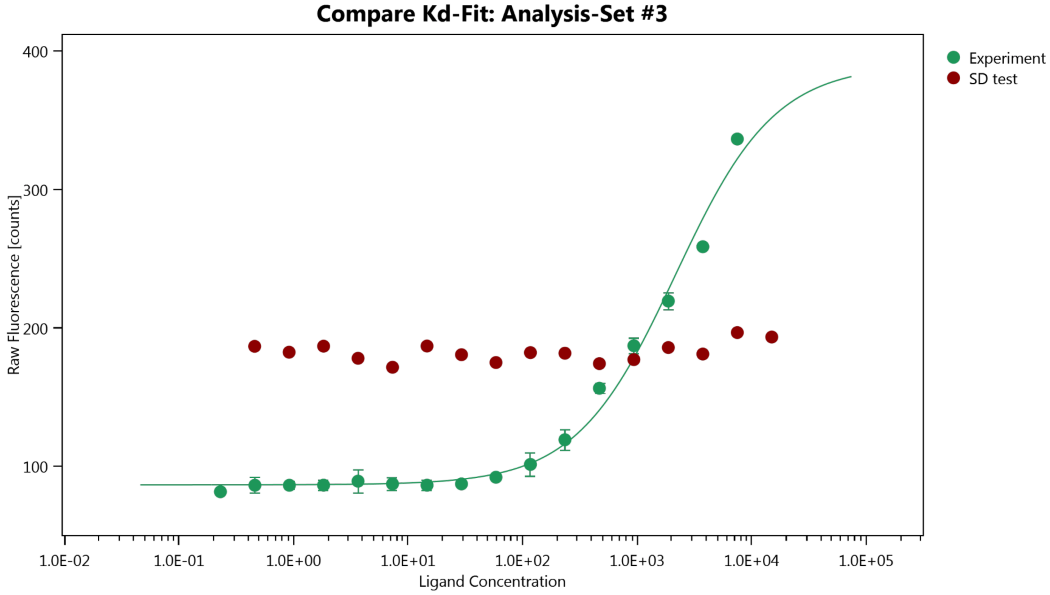

Compare Kd-Fit: Analysis-Set #3
Experiment
SD test
Raw Fluorescence [counts]
100
200
300
400
1.0E-02
1.0E-01
1.0E+00
1.0E+01
1.0E+02
1.0E+03
1.0E+04
1.0E+05
Ligand Concentration

**Supplementary Table S1: Theoretical mass and measured molecular weight, determined by mass photometry of all *Tb*CBCs.**

| Protein complex | Theoretical MW | Measured MW | Events detected |
|---|---|---|---|
| *Tb*CBC-tetramer | 228 kDa | 230 kDa ± 13.2 kDa | 1345 |
| *Tb*CBC-trimer | 162 kDa | 189 kDa ± 27 kDa | 2283 |
| *Tb*CBC dimer | 132 kDa | 135 kDa ± 15.5 kDa | 3066 |
| *Tb*CBC66* | 76 kDa | 81 kDa ± 18.5 kDa | 1684 |

**Supplementary Table S2: Affinity of different protein complexes and RNA as determined by RNA-EMSA**

| Protein complex | RNA | $k_D$ (nM) | sdev |
|---|---|---|---|
| *Tb*CBC-dimer | cap4-SLe | 620 | 46 - 1376 |
| *Tb*CBC-dimer | cap0-SLe | 668 | 395 - 996 |
| *Tb*CBC-dimer | OH-SLe | N/A | N/A |
| *Tb*CBC-trimer | cap4-SLe | 208 | 102 - 320 |
| *Tb*CBC-trimer | cap0-SLe | 179 | 89 - 272 |
| *Tb*CBC-trimer | OH-SLe | 3805 | 2838 - 5548 |
| *Tb*CBC-tetramer | cap4-SLe | 47 | 20 - 67 |
| *Tb*CBC-tetramer | cap0-SLe | 39 | 9 - 61 |
| *Tb*CBC-tetramer | OH-SLe | 135 | 82 - 193 |
| *Tb*CB66* | OH-SLe | 618 | 494 - 760 |
| *Tb*CB66* | 15nt-single stranded | N/A | N/A |
| *Tb*CB66* | SL-hairpin_loop | 390.0 | 296 - 502 |

**Supplementary Table S3: Affinity of *Tb*CBP66 and OH-SLe RNA as determined by fluorescence polarization**

| Protein complex | RNA | $k_D$ (nM) | sdev |
|---|---|---|---|
| *Tb*CB66* WT | OH-SLe | 2545 | 1826 - 3722 |
| *Tb*CB66* R352E | OH-SLe | 2571 | 1955 - 4099 |
| *Tb*CB66* R142E | OH-SLe | N/A | N/A |
| *Tb*CB66* R44E | OH-SLe | N/A | N/A |

**Supplementary Table S4: Gene accession number of *Tb*CBC subunits.**

| Protein name | Gene accession number | UniProt ID | Tag |
|---|---|---|---|
| *Tb*CBP20 | Tb927.6.1970 | Q585L4 | None |
| *Tb*CBP30 | Tb927.10.15210<br>Tb10.61.1300 | Q387Z0 | None |
| *Tb*CBP66 | Tb927.3.1340 | Q57XW7 | None |
| *Tb*CBP110 | Tb927.10.2990<br>Tb10.70.4560 | Q38BU6 | N-Terminal 8xHis |

**Supplementary Table S5. Sequences and characterization data of chemically and chemo-enzymatically synthesized RNAs.**

| RNA | Sequence 5' to 3' | nt | Scale | Isolated yield* | m.w. (calc.) | m.w. (found) |
|---|---|---|---|---|---|---|
| cap0 | Gppp-AACUAACGCUAUUAUUA | 18 | 1 µmol | 42 nmol | 5845.4 | 5845.1 |
| | $m^7$Gppp-AACUAACGCUAUUAUUA | 18 | 30 nmol | 22 nmol | 5859.5 | 5859.2 |
| | $m^7$Gppp-AACUAACGCUAUUAUUA-<br>GAACAGUUUCUGUACUAUAUUG | 40 | 10 nmol | 6 nmol | 12886.6 | 12886.1 |
| cap4 | Gppp-($m^6_2$Am)(Am)(Cm)($m^3$Um)AA-CGCUAUUAUUA | 18 | 1 µmol | 34 nmol | 5943.6 | 5943.7 |
| | $m^7$Gppp-($m^6_2$Am)(Am)(Cm)($m^3$Um)-AACGCUAUUAUUA | 18 | 30 nmol | 20 nmol | 5957.7 | 5957.6 |
| | $m^7$Gppp-($m^6_2$Am)(Am)(Cm)($m^3$Um)-<br>AACGCUAUUAUUAGAACAGUUU-<br>CUGUACUAUAUUG | 40 | 10 nmol | 5 nmol | 12984.8 | 12985.0 |

*Amount of pure product obtained after HPLC purification
